# Efficient whole genome sequencing of influenza A viruses

**DOI:** 10.1101/749234

**Authors:** Marina Escalera-Zamudio, Ana Georgina Cobián-Güemes, Blanca Taboada, Irma López-Martínez, Joel Armando Vázquez-Pérez, Maricela Montalvo-Corral, Jesus Hernandez, José Alberto Díaz-Quiñonez, Gisela Barrera Badillo, Susana López, Carlos F. Arias, Pavel Iša

## Abstract

The constant threat of emergence for novel pathogenic influenza A viruses with pandemic potential, makes full-genome characterization of circulating influenza viral strains a high priority, allowing detection of novel and re-assorting variants. Sequencing the full-length genome of influenza A virus traditionally required multiple amplification rounds, followed by the subsequent sequencing of individual PCR products. The introduction of high-throughput sequencing technologies has made whole genome sequencing easier and faster. We present a simple protocol to obtain whole genome sequences of hypothetically any influenza A virus, even with low quantities of starting genetic material. The complete genomes of influenza A viruses of different subtypes and from distinct sources (clinical samples of pdmH1N1, tissue culture-adapted H3N2 viruses, or avian influenza viruses from cloacal swabs) were amplified with a single multisegment reverse transcription-PCR reaction and sequenced using Illumina sequencing platform. Samples with low quantity of genetic material after initial PCR amplification were re-amplified by an additional PCR using random primers. Whole genome sequencing was successful for 66% of the samples, whilst the most relevant genome segments for epidemiological surveillance (corresponding to the hemagglutinin and neuraminidase) were sequenced with at least 93% coverage (and a minimum 10x) for 98% of the samples. Low coverage for some samples is likely due to an initial low viral RNA concentration in the original sample. The proposed methodology is especially suitable for sequencing a large number of samples, when genetic data is urgently required for strains characterization, and may also be useful for variant analysis.

## INTRODUCTION

Influenza A viruses (IAVs) infect a wide range of avian and mammalian species, including humans. IAVs are an important cause of human respiratory diseases, generating seasonal yearly infections and occasional pandemics^1^. These enveloped RNA viruses belong to the Orthomyxoviridae family, having a segmented genome composed of eight independent RNA segments, ranging in size from 890 to 2340 nucleotides. The eight different genome segments encode for over ten structural and no-structural viral proteins: PB2, PB1 and PA (forming the polymerase complex), HA (hemagglutinin protein, responsible for binding to cell receptors and is the main antiviral target), NP (nucleoprotein), NA (neuraminidase protein, that cleaves sialic acid allowing virion liberation, and is also an antiviral target), M1 and M2 (matrix and viral ion channel proteins) and NS1 and NS2 (non-structural proteins, major host immune modulators). Other nonessential accessory proteins, with a wide variety of functions, include PB1-F2, PB1-N40, PA-X, PA-N182 and PA-N155^2^. Based on the variability of the two surface glycoproteins (HA and NA), 18 HA and 11 NA varieties have been described so far, that combined generate different virus subtypes^3–5^. While almost all viral subtypes are believed to circulate in waterfowl, only three IAVs subtypes have been established in human populations (H1N1, H2N2, and H3N2)^1^.

IAVs evolve mainly by two mechanisms: single point mutations introduced into different genes by a low-fidelity RNA polymerase that may be fixed under varying selective pressures (antigenic drift), or by the exchange of whole segments between different viral strains during co-infection (antigenic shift)^1^. Both of these processes contribute to the long-term evolution of IAVs, and are associated with changes in antigenic or biological properties of emerging strains (such as an increased virulence, change of host tropism, resistance to antiviral drugs and antigenic escape, among others)^6–8^.

The evolution of circulating IAVs has been mainly studied through the HA and NA genes^9^, as whole genome sequences only became widely available within the last decade, after the introduction of the high-throughput sequencing (HTS) technologies ^10–12^. Before HTS, the most common method for obtaining complete IAVs genomes was by the amplification of overlapping genome regions using a reverse transcription polymerase chain reaction (RT-PCR), followed by Big Dye Terminator sequencing chemistry^10,12–14^. Since then, HTS has been successfully used to obtain whole genome sequences directly from clinical samples, after cell culture adaptation, or directly after RT-PCR amplification of individual genome segments (namely HA and NA)^15–20^. Multisegment reverse transcription-PCR reaction (M-RTPCR) is often used to amplify the genetic material to be used for preparation of sequencing libraries^21–24^. However, the quality and coverage of the viral sequences obtained can vary greatly, ranging from a high genome coverage with > 100x depth, to only a few reads per sample^15,18,20–24^. We present here a simple and cost-efficient methodology to obtain whole genome sequence from IAVs with a good success rate, even when started from a non-optimal amount of initial genetic material.

## MATERIAL AND METHODS

### Samples collection and viral RNA extraction

The Bioethics Committee of Instituto de Biotecnologia UNAM, the INER (Instituto Nacional de Enfermedades Respiratorias), and the Hospital Juarez approved human samples collection and all handling protocols used in this work. All hospitalized patients read, agreed and signed to written consent forms. The twenty-nine viral samples from different sources used for this work are listed in Table 1. For tissue culture-adapted H3N2 viruses and for avian influenza A viruses collected from cloacal swabs, 200 μl of the supernatants were treated with Turbo DNase (Ambion) for 30 minutes at 37°C to deplete foreign DNA. Viral RNA was then extracted with the Purelink Viral RNA/DNA extraction kit (Invitrogen), using 25 μg of linear acrylamide (Ambion) as a carrier. RNA was then stored at -70°C for further use. For clinical H1N1 samples, viral RNA was extracted from 200 μl of the sample using a high-throughput system (MagNA Pure®, Roche, Indianapolis, IN, USA), as described by the manufacturer. We did not observe important differences among samples, as both high and low amounts of genomic material were obtained using both extraction systems as quantified by qPCR (data not shown), indicating that the different extraction processes do not affect final results, and that the genomic material quality and quantity is highly dependent on the quality of the initial sample.

**Table 1.** Whole genome sequencing of AIVs.

| Sample | Source<sup>a</sup> | Method<sup>b</sup> | Starting material (ng)<sup>c</sup> | Total reads<sup>d</sup> | Unique reads<sup>e</sup> | Unique/Total(%)<sup>f</sup> | IAV-specific reads<sup>g</sup> | Normalized Reads<sup>h</sup> | Genome coverage (nt, %)<sup>k</sup> | Depth<sup>l</sup> | HA coverage (nt, %)<sup>m</sup> | NA coverage (nt, %)<sup>m</sup> |
| --- | --- | --- | --- | --- | --- | --- | --- | --- | --- | --- | --- | --- |
| A/Mexico/InDRE835/2003 (H3N2) | CC | FR | 2000 | 5,516,940 | 694,502 | 12.6 | 390,379 | 56.2 | 13,627 (100) | > 50x | 1,762 (100) | 1,467 (100) |
| A/Mexico/InDRE2112/2005 (H3N2) | CC | FR | 2600 | 7,968,740 | 448,404 | 5.6 | 195,195 | 43.5 | 13,609 (99.9) | >50x | 1,760 (99.9) | 1,467 (100) |
| A/Mexico/InDRE2160/2004 (H3N2) | CC | FR | 2700 | 6,242,192 | 841,094 | 13.5 | 394,596 | 46.9 | 13,618 (99.9) | >50x | 1,756 (99.7) | 1,467 (100) |
| A/Mexico/InDRE2601/2005 (H3N2) | CC | FR | 2300 | 3,200,194 | 299,053 | 9.3 | 124,813 | 41.7 | 13,609 (99.9) | >50x | 1,760 (99.9) | 1,467 (100) |
| A/Mexico/InDRE2662/2003 (H3N2) | CC | FR | 2400 | 8,396,552 | 617,408 | 10.9 | 133,250 | 21.6 | 13,624 (99.9) | >50x | 1,762 (100) | 1,466 (99.9) |
| A/Mexico/IBT22/2009 (H1N1) | swab | FR | 900 | 5,718,955 | 725,937 | 12.7 | 100,944 | 13.9 | 13,630 (99.9) | 30X | 1,777 (100) | 1,457 (99.9) |
| A/Mexico/IBT23/2009 (H1N1) | swab | FR | 1400 | 6,406,144 | 861,913 | 13.5 | 183,632 | 21.3 | 12,835 (94.2) | 25X | 1,776 (99.9) | 1,458 (100) |
| A/Mexico/INER2/2009 (H1N1) | swab | FR | 500 | 2,966,105 | 649,594 | 21.9 | 16,091 | 2.5 | 12,971 (95.2) | 20X | 1,773 (99.8) | 1,455 (99.8) |
| A/Mexico/INER11/2010 (H1N1) | swab | FR | 600 | 5,498,044 | 380,564 | 6.9 | 182,769 | 48.0 | 13,632 (100) | > 50x | 1,777 (100) | 1,458 (100) |
| A/green winged teal/Sonora/266/2008 (H6N1) | swab | FR | 610 | 8,990,438 | 813,130 | 9 | 122,616 | 15.0 | 13,600 (99.8) | > 50x | 1,737 (100) | 1,458 (100) |
| A/redhead/Sonora/408/2008 (H5N2) | swab | FR | 430 | 7,011,978 | 834,433 | 11.9 | 235,457 | 28.2 | 11,743 (86.7) | 25X | 1,767 (100) | 1,020 (70.2) |
| A/green winged teal/Sonora/701/2008 (H11N3) | swab | FR | 800 | 7,561,429 | 832,664 | 11 | 224,200 | 26.9 | 13,583 (99.7) | > 50x | 1,760 (100) | 1,453 (100) |
| A/northern shoveler/Sonora/738/2008 (H6N1) | swab | FR | 750 | 6,081,579 | 503,531 | 8.3 | 302,830 | 60.1 | 13,613 (99.9) | > 50x | 1,747 (100) | 1,458 (100) |
| A/green winged teal/Sonora/1116/2009 (H10N7) | swab | FR | 450 | 5,260,599 | 548,112 | 10.4 | 382,955 | 69.9 | 13,618 (99.9) | > 50x | 1,728 (100) | 1,461 (100) |
| A/Mexico/InDRE756/2003 (H3N2) | CC | PCR | 54 | 89,978 | 29,563 | 32.9 | 6,218 | 21.0 | 13,581 (99.7) | 8X | 1,762 (100) | 1,467 (100) |
| A/Mexico/InDRE2227/2005 (H3N2) | CC | PCR | 10 | 6,035,458 | 787,564 | 13 | 229,130 | 29.1 | 13,624 (99.9) | >50x | 1,762 (100) | 1,467 (100) |
| A/Mexico/InDRE2246/2005 (H3N2) | CC | PCR | 120 | 18,937,312 | 2,113,683 | 11.2 | 389,053 | 18.4 | 13,627 (100) | >50x | 1,762 (100) | 1,467 (100) |
| A/Mexico/INER9/2010 (H1N1) | swab | PCR | 13 | 7,618,995 | 1,599,105 | 21 | 4,115 | 5.8 | 10,685 (78.4) | 30X | 1,747 (98.3) | 1,446 (99.2) |
| A/Mexico/INER10/2009 (H1N1) | swab | PCR | 28 | 5,495,437 | 1,591,161 | 29 | 17,496 | 11.0 | 12,290 (90.2) | 10X | 1,774 (99.8) | 1,448 (99.3) |
| A/Mexico/INER12/2009 (H1N1) | swab | PCR | 30 | 1,145,281 | 167,785 | 14.7 | 32,437 | 19.3 | 12,218 (89.6) | 15X | 1,777 (100) | 1,458 (100) |
| A/Mexico/INER13/2009 (H1N1) | swab | PCR | 116 | 4,813,830 | 670,629 | 13.9 | 16,703 | 2.5 | 11,100 (81.4) | 10X | 1,687 (95.5) | 1,361 (93.3) |
| A/Mexico/INER14/2009 (H1N1) | swab | PCR | 116 | 4,265,616 | 1,001,792 | 23.5 | 222,467 | 22.2 | 13,631 (99.9) | > 50x | 1,776 (99.9) | 1,458 (100) |
| A/Mexico/INER15/2009 (H1N1) | swab | PCR | 41 | 2,464,905 | 439,251 | 17.8 | 4,783 | 1.1 | 9,207 (67.5) | 30X | 1,666 (93.4) | 1,360 (93.3) |
| A/Mexico/INER17/2009 (H1N1) | swab | PCR | 51 | 5,056,489 | 1,304,019 | 25.8 | 34,615 | 2.7 | 10,988 (80.6) | 10X | 1,695 (95.4) | 1,435 (98.4) |
| A/Mexico/INER18/2010 (H1N1) | swab | PCR | 40 | 5,444,679 | 1,021,111 | 18.8 | 42,142 | 4.1 | 10,245 (75.2) | 10X | 1,772 (99.7) | 1,458 (100) |
| A/American wigeon/Sonora/769/2008 (H9N2) | swab | PCR | 115 | 10,008,629 | 1,910,239 | 19 | 580,907 | 30.4 | 13,627 (100) | > 50x | 1,742 (100) | 1,409 (97) |
| A/northern shoveler/Sonora/797/2008 (H5N3) | swab | PCR | 130 | 10,902,300 | 2,589,322 | 23.8 | 582,465 | 22.5 | 13,626 (100) | > 50x | 1,773 (100) | 1,453 (100) |
| A/green winged teal/Sonora/829/2009 (H6N5) | swab | PCR | 135 | 7,855,551 | 1,928,906 | 24.6 | 310,219 | 16.1 | 13,627 (100) | > 50x | 1,738 (100) | 1,469 (100) |
| A/green winged teal/Sonora/1132/2009 (H10N3) | swab | PCR | 90 | 7,318,431 | 2,106,937 | 28.8 | 192,033 | 9.1 | 13,489 (99) | > 50x | 1,728 (100) | 1,453 (100) |

a CC: cell-culture derived viruses grown in Madin-Darby Canine Kidney (MDCK). For clinical samples, nasopharyngeal swabs were obtained. For avian samples, cloacal swabs were obtained.
b Method used for sample processing. FR stands for fragmentase digestion after M-RT-PCR. PCR stands for an additional PCR amplification using random primers, with no fragmentation step.
c Amount of starting material in ng obtained after M-RT-PCR amplification.
d Number of quality-filtered reads obtained per sample after sequencing.
e Number of unique reads after collapsing identical reads.
f Percent of unique reads out of total reads.
g Number of influenza-specific unique reads.
h Percent of influenza-specific reads out of unique reads.
k Genome coverage in base pairs, whilst the corresponding percentage is shown in brackets.

### Whole genome amplification

Complete genomes for all IAV samples were amplified with a single multisegment reverse transcription-PCR reaction (M-RTPCR) as described by Zhou et al ^25^, using 5 μl of extracted RNA as input for each reaction. PCR products were visualized on 1% agarose gels and then purified using firstly by the AMPure magnetic beads (Beckman Coulter Genomics), and later by the DNA Clean & Concentrator kit (Zymo Research). All final products were quantified using the NanoDrop ND1000 (NanoDrop Technologies) and the Agilent 2100 Bioanalyzer.

### Preparation of Illumina libraries and sequencing

Samples with more than 150ng of amplified material after M-RTPCR were subjected directly to the preparation of HTS libraries, as described below. DNA was fragmented to approximately 250-500 bp using the NEBNext Fragmentase enzyme (New England Biolabs), and further purified with the DNA Clean & Concentrator kit (Zymo Research). Products were used for building individual single-indexed Illumina libraries using the Genomic DNA Sample Preparation kit - Multiplex Sample Prep Oligo kit (Illumina), as instructed by the manufacturer. At the time, Illumina protocols recommended between 1 and 5 μg of DNA as starting material. Thus, when less than 150ng of total genomic material was obtained after initial M-RTPCR, these samples were additionally amplified using a PCR with random primers, as described in Taboada et al ^26^. In this protocol, no additional enzymatic fragmentation step is needed, given the nature of the reaction. Briefly, two rounds of synthesis with Sequenase 2.0 (USB, USA) were performed using primer A (5’-GTTTCCCAGTAGGTCTCN-3’), followed by ten amplification rounds using the Phusion DNA polymerase (Finnzymes) with primer B (5’-GTTTCCCAGTAGGTCTC-3’). DNA was then digested using the *GsuI* enzyme to remove the 16 additional nucleotides from 5’ and 3’ ends generated during the PCR reaction. Digested DNA was purified as described above, and then used to prepare Illumina sequencing libraries, as described above.

Between 4 to 6 libraries of approximately 350 bp were equimolarly pooled and loaded per lane to generate sequencing clusters, followed by 36 or 45 cycles of single base pair extensions using the Illumina Genome Analyzer II platform at the HTS Core Facility-UNAM. Image analysis was done using the Genome Analyzer Pipeline Version 1.4 (Illumina, San Diego, CA). As a negative control, the pure distilled water was included from the initial M-RTPCR and further processes as a sample for the preparation of a blank library (data not shown).

### Bioinformatic analysis of sequencing data

Data analysis was performed using the computational cluster of the Instituto de Biotecnologia-UNAM, as described in Escalera et al ^27,28^. High-quality reads (Q30) were filtered and identical reads were collapsed. The reference genomes of the A/Netherlands/602/2009 (H1N1) and A/New York/392/2004 (H3N2) strains were used to filter out H1N1 and H3N2 viral sequences, using the mapping and assembly software MAQ v0.7.1 ^29^. For viruses of avian origin, the reads obtained per sample were first mapped against an *in-house* curated database, as described by Paulin et al ^30^. The reference strains with most reads mapped were selected for further genome assembly. Mapping of reads to the reference genomes was performed using SMALT v0.7.1, whilst consensus sequences were called using SAMtools version 0.1.18^31^. Given that avian IAVs genomes could have a higher variability when compared to human H1N1 or H3N2 samples, two separate mapping rounds were performed. In the first round, 15 mismatches were allowed to generate a ‘relaxed’ consensus sequence, whilst during the second round only 5 mismatches were allowed for mapping to allow for a ‘strict’ consensus using as a reference the consensus sequence generated during the first round. Reads that did not mapped during the second round were not considered for further analysis. Finally, influenza-specific reads were used to plot genome coverages, using the R package. The sequences generated in this work are available in GenBank under the following accession numbers: CY100445-CY100622, KY575169-KY575224, KY593192-KY593200.

### Statistical analysis

The differences in *i)* the ratio of total to unique reads (after collapsing identical), *ii)* in the ratio of unique to influenza-specific reads, and *iii)* in the mean of total normalized reads as a function of the sample processing method (M-RTPCR/enzymatic fragmentation [FR] vs. additional PCR amplification with random primers and no enzymatic fragmentation [PCR]), were statistically assessed under an unpaired (two sample means) two-tailed t-test in Prism 6.0. Given the sample size for each group (n=14 for FR and n=15 for PCR) and the mean and standard deviation values for each class, a statistical power of >80% was estimated with a Type 1 error rate of 5%.

## RESULTS AND DISCUSSION

### Viral RNA enrichment

The M-RTPCR described by Zhou et al ^25^ is based on all IAVs having conserved 5’ and 3’ ends for all genome segments. Thus, all segments can be amplified using the same pair of oligonucleotides in a single multiplex reaction. Given the diverse origin of the samples used in this study (culture-grown virus, clinical isolates and avian-derived samples), variable amounts of PCR-amplified whole genome products were obtained, ranging from 10 ng up to 2.7 μg of total DNA/sample (Table 1). When comparing the differential amplification for each genome segment, PCR products corresponding to the smaller segments (M and NS, followed by HA, NP and NA) were favored against the larger ones (PB2, PB1, PA). In general, the efficiency (yield) of PCR was greater for low-molecular than for a high-molecular weight DNA templates (Figure 1). For 15 samples, less than 150 ng of total genomic DNA was obtained (Table 1), thus these were subjected to a secondary random PCR amplification as described in material and methods.

**Figure 1.**
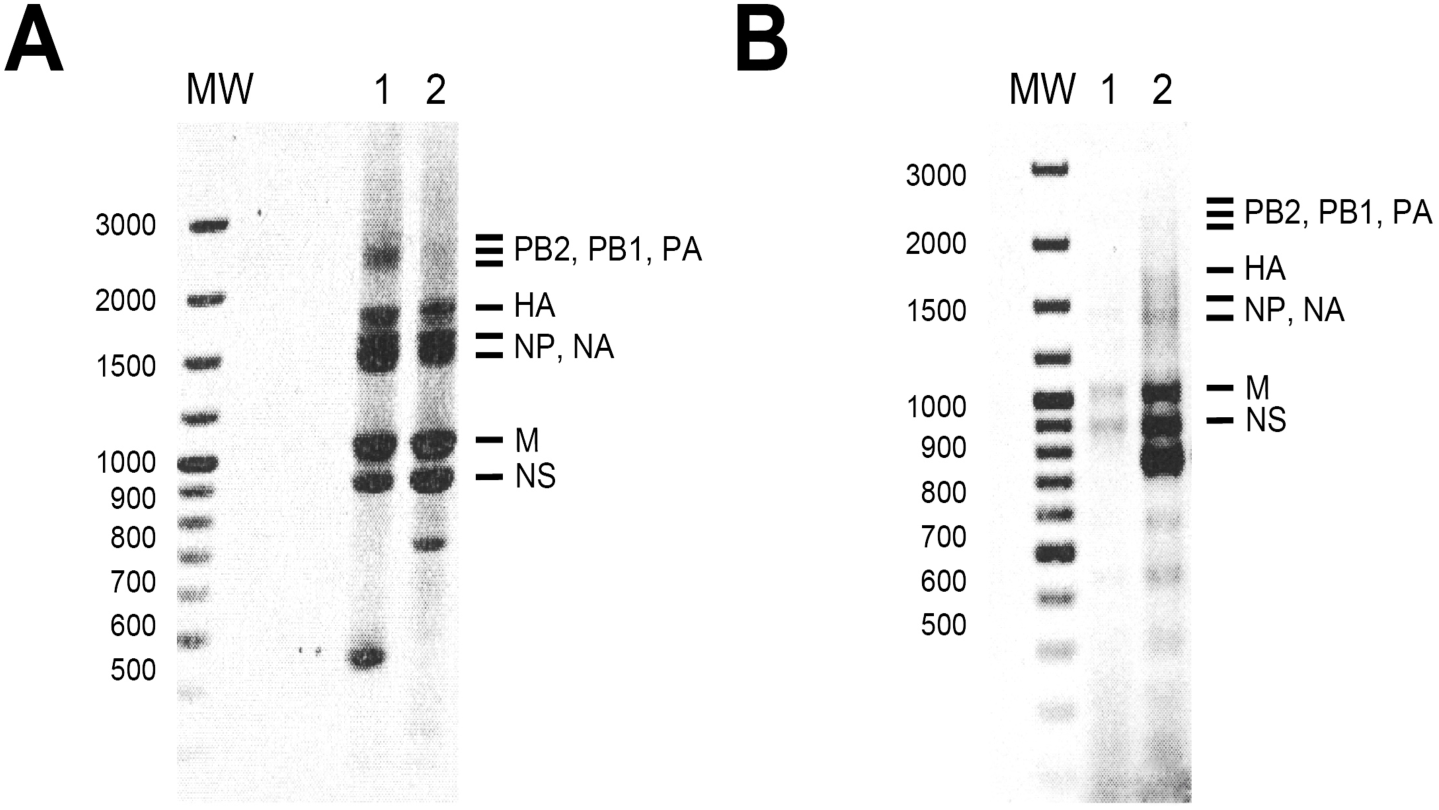
Influenza A virus genome amplification by M-RTPCR. Influenza A virus samples of different origins were amplified using a multisegment reverse transcription-PCR reaction (M-RTPCR), and visualized on a 1% agarose gel. Molecular weight markers are indicated on the left side. A. Amplification of human influenza A strains, lane 1: A/Mexico/InDRE2662/2003 (H3N2); lane 2: A/Mexico/IBT23/2009 (H1N1). B. Amplification of avian samples, lane 1: A/green winged teal/Sonora/701/2008 (H11N3); lane 2: A/northern shoveler/Sonora/738/2008 (H6N1).

### Sequencing specificity

In total, 29 IAV genomes were sequenced, with a total number of reads per sample ranging from 90,000 to 19×10^6^ (Table 1). The lowest number of raw reads was obtained for sample A/Mexico/InDRE756/2006 (H3N2), whilst the highest one was for sample A/Mexico/InDRE2246/2005 (H3N2). Given the low concentration of initially amplified genomic products obtained after M-RTPCR, these two samples were subjected to additional amplification by random PCR (Table 1). Identical reads were collapsed, reducing the number of reads by a mean of 84% (ranging from 70% to 95%). When comparing the mean percentage of unique reads with respect to the method of library preparation, a significant difference (p-value <0.0001) was observed between samples prepared by the FR method (mean: 11.25 ± 3.9), and samples processed under the PCR method (mean: 21.19 ± 6.5), with PCR method having higher proportion of unique reads (Figure 2A). When comparing the number of influenza-specific reads obtained under the different methods, samples processed under the PCR method showed a significantly smaller number of influenza-specific reads (14.7 ± 2.7, p-value < 0.01), when compared to samples prepared by the FR method (30.3 ± 5.1) (Figure 2B). This observation suggests that despite yielding a reduced number of unique reads after sequencing, the FR method increased specificity (as shown by a larger number of influenza-specific reads), rendering it better method when compared to the PCR method.

**Figure 2.**
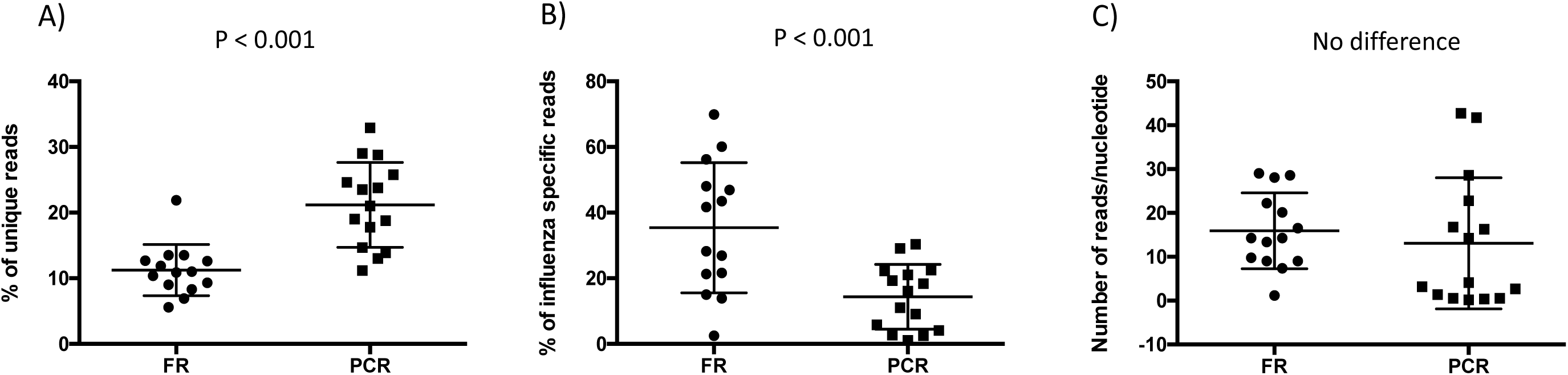
Differences in the number of reads as a function of the sample processing methods. A. The mean percentage of unique reads (after collapsing identical) obtained per sample under different processing methods (FR: M-RTPCR/enzymatic fragmentation, PCR: additional PCR amplification using random primers and no enzymatic fragmentation). Differences within means were statistically assessed under an unpaired (two sample means) two-tailed t-test (p-values shown). B. The mean percentage of influenza-specific reads (determined after mapping to reference genomes, numbers are normalized from the total number of unique reads per sample). Differences within means were statistically assessed under an unpaired (two sample means) two-tailed t-test (p-values shown). C. Number of influenza-specific reads per sequenced nucleotide of IAV genome.

Additionally, when the number of influenza-specific reads was compared to genome coverage, samples prepared under the FR method showed a larger number of reads per nucleotide sequenced, as the majority of samples displayed between 10-20 reads per nucleotide. Contrastingly, >50% of the samples prepared by the PCR method showed < 5 reads per nucleotide sequenced (Figure 2C). A lower percentage of influenza specific reads obtained under the PCR method could be partially explained by non-specific amplification of non-viral genetic material during the random PCR. Thus, it is important to note that too many rounds of additional amplification by random PCR may introduce sequencing bias, and can also increase the relative quantities of non-viral genetic material. However, the PCR method does facilitate whole genome sequencing when samples have an initial low concentration of genomic material after M-RTPCR product (Table 1).

### Genome coverage

IAV whole genome sequencing studies have shown variable results, as obtaining full genomic sequences can be difficult, especially for clinical or tissue-derived samples^15–20^. Whole genome sequencing was highly efficient, with > 99% genome coverage obtained for 66% of the samples (19/29): 11 (79%) for FR method and 8 (53%) for PCR. Many samples (59%, 17/29) also showed good overall sequencing depth across the whole genome (>50x) (Table 1). For the majority of the samples (20/29), more than 100,000 unique influenza-specific reads were obtained, supporting for the methodology proposed here as suitable for detection viral variants within a single sample. For some samples, gaps were observed within genome assemblies, mainly within the large genome segments (namely in PB2, PB1 and PA) (Figure 3). This can be explained to some extent by the M-RTPCR bias towards the amplification of smaller segments. Incomplete genome sequencing is also correlated to low viral RNA concentrations, in relation to the quality of the original sample.

**Figure 3.**
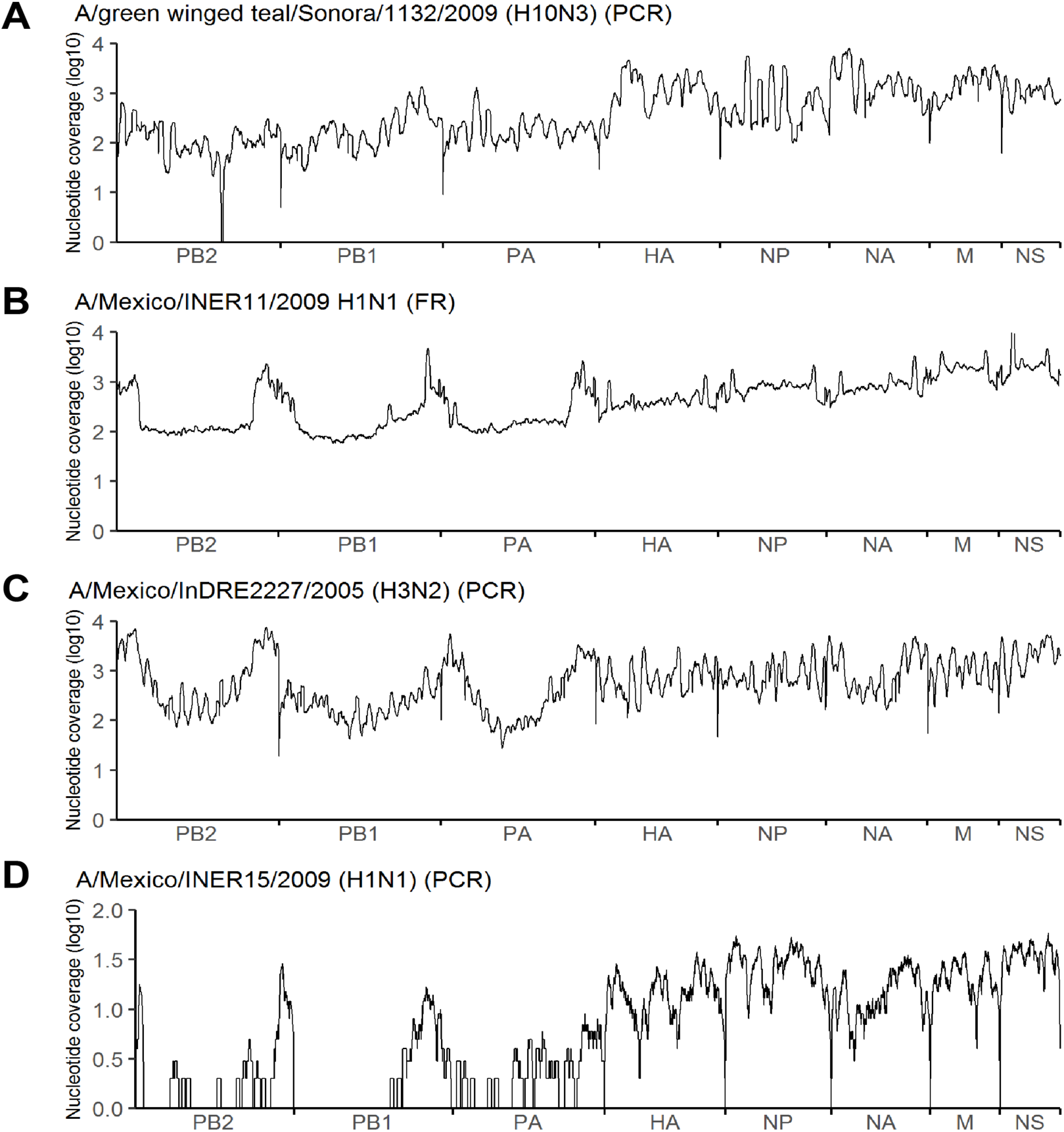
Whole genome coverage for representative virus samples sequenced. For each sample, genome coverage is plotted on a log-scale, as a function of the genome nucleotide position. The corresponding viral genome segments are shown on the X axis. Four different representative samples were used for visualization example. A. A/green winged teal/Sonora/1132/2009 (H10N3), (prepared under the PCR method, see text for description); B. A/Mexico/INER11/2009 H1N1 (prepared under the FR method); C. A/Mexico/InDRE2227/2005 H3N2 (prepared under the PCR method) and D. A/Mexico/INER15/2009 H1N1 (prepared under the PCR method

Nonetheless, even in samples with lower genome coverage, the antigenically relevant segments (HA and NA), which are of mayor importance for epidemiological surveillance, were fully sequenced for 79% of the samples (23/29), with over 99.3% coverage and a minimum depth of 10X (Table 1). Samples with good quality and quantity of initial genomic material after M-RTPCR showed an overall good genome coverage and depth (Table 1). Comparably, samples with a low amount of starting genomic material after M-RTPCR (and thus reamplified by random PCR), also showed good genome coverage and depth, confirming the usefulness of both methods for obtaining whole genome sequences.

## CONCLUSIONS

In this work, we present a simple methodology to improve the likelihood of obtaining whole genome sequences with good genome coverage and depth, for non-optimal samples. This methodology is especially suitable for sequencing a large number of samples, and is useful when genetic data is needed to perform evolutionary or epidemiological analyses during outbreaks. The proposed methodology also allows the identification and characterization of field samples difficult to sequence by conventional molecular biology methods. This methodology is also compatible with improved library preparation kits and sequencing technologies, such as the MinION portable device, that require less initial genomic material.

## ACKNOWLEDGMENT

We thank Dr. Ricardo Grande and M.Sc. Verónica Jiménez and the HTS core facility of the National University of Mexico for their technical support. Computational analysis was performed using the cluster of the Instituto de Biotecnología-UNAM. We thank Dr. Daniel Aguilar Angeles from the Hospital Juárez de México for facilitating clinical samples. This work was supported by grants 5549 and I0110/184/09 from the National Council for Science and Technology CONACyT-Mexico, and grant PICOSI09-209 from the Instituto de Ciencia y Tecnologia del Distrito Federal. Marina Escalera-Zamudio and Georgina Cobián-Güemes were supported by a scholarship from CONACyT-Mexico. Marina Escalera-Zamudio is currently supported by an EMBO Long Term Fellowship (ALTF376-2017).

